# Mycobacteria-host interactions in human bronchiolar airway organoids

**DOI:** 10.1101/2020.11.12.379586

**Authors:** Nino Iakobachvili, Stephen Adonai Leon Icaza, Kèvin Knoops, Norman Sachs, Serge Mazères, Roxane Simeone, Antonio Peixoto, Marlène Murris-Espin, Julien Mazières, Carmen López-Iglesias, Raimond B.G. Ravelli, Olivier Neyrolles, Etienne Meunier, Geanncarlo Lugo-Villarino, Hans Clevers, Céline Cougoule, Peter J. Peters

## Abstract

Tuberculosis, one of the oldest human pathogens remains a major global health threat. Recent advances in organoid technology offer a unique opportunity to grow different human “organs” *in vitro*, including the human airway, that faithfully recapitulate tissue architecture and function. We have explored the potential of human airway organoids (AOs) as a novel system in which to model tuberculosis infection. To this end, we adapted biosafety containment level 3–approved procedures to allow successful microinjection of *Mycobacterium tuberculosis*, the causative agent of tuberculosis, into AOs. We reveal that mycobacteria infected epithelial cells with low efficiency, and that the organoid microenvironment was able to control, but not eliminate the pathogen. We demonstrate that AOs responded to infection by inducing cytokine and antimicrobial peptide production, and inhibiting mucins. Given the importance of myeloid cells in tuberculosis infection, we co-cultured mycobacteria-infected organoids with human monocyte-derived macrophages, and found that these cells were recruited to the organoid epithelium. We conclude that adult stem cell–derived airway organoids can be used to model early events of tuberculosis infection and offer new avenues for fundamental and therapeutic research.

## Introduction

Airborne pathogens are a major cause of death worldwide. Respiratory infectious diseases cause more than 5 million fatalities annually, with tuberculosis (TB) accounting for one-fifth (WHO Global tuberculosis report 2019). In 2018, TB caused an estimated 1.5 million deaths, making TB one of the top 10 killers worldwide, and 25% of the worlds population is thought to be latently infected by *Mycobacterium tuberculosis* (Mtb) (1).

The lung is the entry port for Mtb and the main site of TB disease manifestation. Mtb-containing droplets navigate through the lung anatomy and airway functions in order for mycobacteria to establish its replicative niche in the alveolar space (2, 3). Models of human lung infection are therefore crucial to increase our understanding of host–pathogen interactions-an essential step towards new drug development. Whilst conventional 2D cell culture and animal models have contributed to deciphering key host–pathogen mechanisms at play during Mtb infection (4), they lack relevance with the human lung.

One of the major breakthroughs in the stem cell field is the ability to grow human ‘organs’ *in vitro*, also known as organoids (5). Human airway organoids (AOs) are derived from adult stem-cells and composed of a polarized, pseudostratified airway epithelium containing basal, secretory and multi-ciliated cells, although they are currently lacking alveolar pneumocytes. They display functional mucus secretion and ciliate beating (6), therefore constituting a suitable human system in which to model early steps of host–pathogen interactions (7–9). We have set out to evaluate the potential of AOs as a model in which to study Mtb infection. Our data demonstrate that mycobacteria can be readily found in the lumen of AOs with some internalization by airway epithelial cells and overall control of mycobacterial growth. In response to Mtb infection, we show AOs inducing the secretion of cytokines and antimicrobial peptides, and the option to model innate cell recruitment by co-culturing human macrophages with injected AOs.

## Results & Discussion

Due to the innate cystic structure of AOs, where the pathogen-sensing apical side faces the lumen, DsRed-expressing H37Rv Mtb (mean 4271±834 CFU/organoid) was microinjected via a BSL-3–approved custom-made micro-injection system (Figure 1A). Bacteria could be found in the lumen of AOs and occasionally making contact with epithelial cells but without causing obvious alterations to organoid architecture and ultrastructure (Figure 1B-C, Movie S1 and S2). Evident from the movies is the functional mucociliary system where cilia beat secreted mucus and cell debris around the lumen.

**Figure 1.**
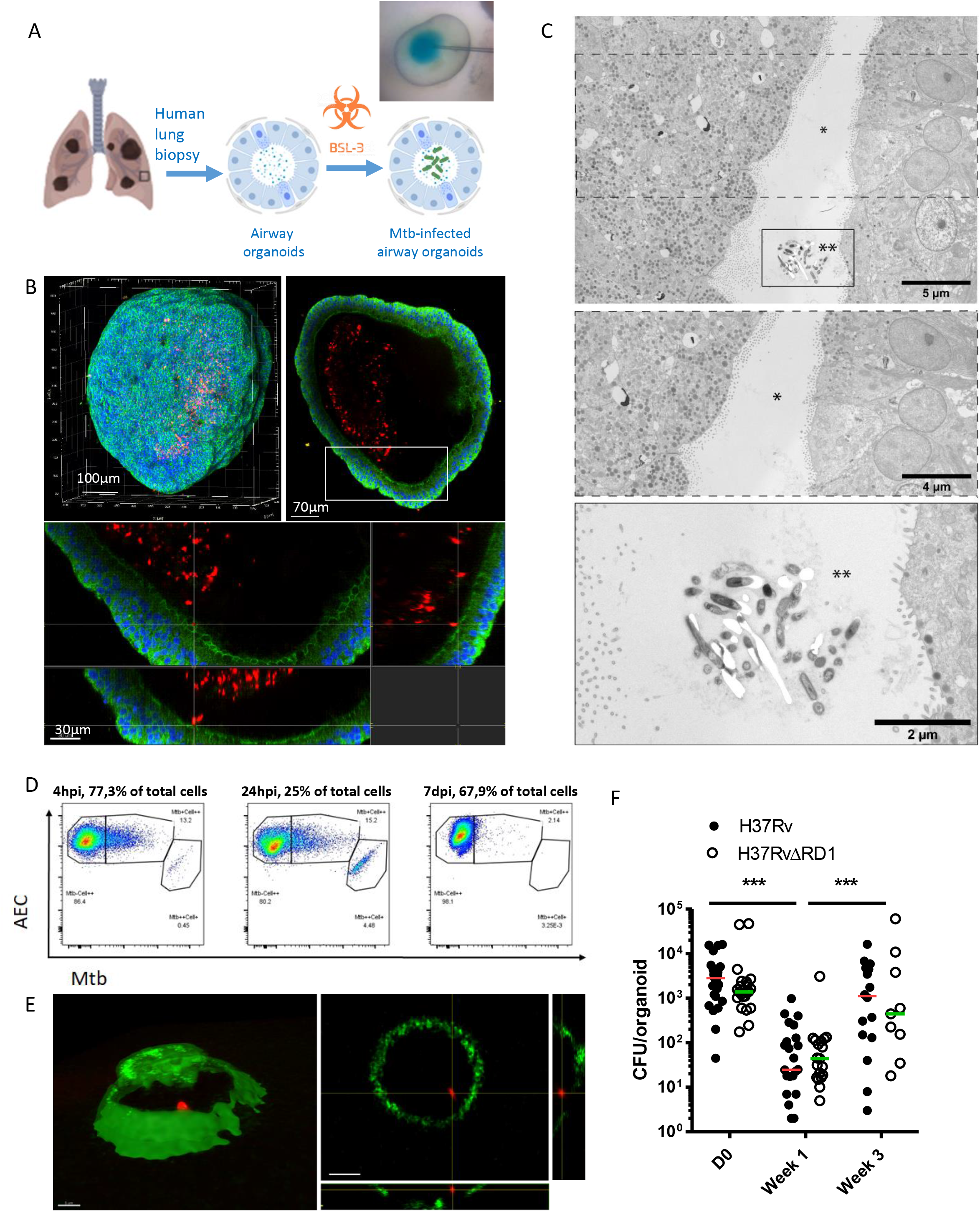
Human airway organoids (AOs) infected with Mtb. (A) Experimental scheme and bright-field image of the microinjection. (B) Confocal microscopy of DsRed-expressing H37Rv inside AOs, 4 days post-infection. Nuclei are labeled with DAPI (blue); cellular membranes with CellMask green (green). (C) Transmission electron microscopy at one week post-infection showing H37Rv within the organoid lumen. Lower panels show magnifications of the boxed areas in the upper image. (D) Quantification of cells associated with H37Rv after 4 (left) or 24 hours (middle) incubation with AOs-derived single cells or 7 days incubation in whole organoids (right) (E) Representative images of sorted epithelial cells with intracellular DsRed-expressing H37Rv, scale bars = 5μm. (F) CFU counts from individual organoids on the day of microinjection (day 0), 7 and 21 days post-infection. Each dot represents one organoid. Lines indicate median CFU counts. The experiment was performed at least four times independently. ***P < 0.001 by a two-tailed Mann-Whitney test.

Mtb is known to infect bronchial epithelial cells in 2D conditions (10), and pneumocytes *in vitro* (11) and *in vivo* (12), but with low efficiency. To identify if Mtb could infect organoid derived epithelial cells, AOs were dissociated into single cells, infected with Mtb H37Rv and analyzed by flow cytometry. Approximately 13% and 19% of epithelial cells were found associated with bacteria after 4 h and 24 h of infection, respectively (Figure 1D). Sorted epithelial cells showed that individual cells harboured Mtb (Figure 1E) suggesting cell invasion by a yet unknown mechanism. The number of internalised bacteria dropped to 2% when AOs, which had been infected with Mtb for 7 days, were dissociated into single cells and analyzed by flow cytometry (Figure 1D). The functioning mucociliary clearance system within AOs is likely responsible for reducing mycobacterial contact with epithelial cells.

Mtb has a functional type VII secretion system (ESX-1) encoded by the RD1 locus which is involved in modulating host responses and inducing host cell lysis (13–16). To determine whether the presence of ESX-1 induced increased epithelial cell lysis, we quantified cell death by TOPRO-3 incorporation in AOs after injection of wild-type H37Rv or H37RvΔESX-1 which lacks ESX-1. Neither strain induced significant epithelial cell death in Mtb-infected AOs compared to uninfected ones (Supp. Figure 1A), indicating that a functional ESX-1 expression does not trigger increased epithelial cell damage.

Next, we investigated mycobacterial survival in AOs. Mtb H37Rv demonstrated a bi-phasic curve (Figure 1F), with a significant decrease of bacterial load after 7 days followed by an increase at 21 days post-infection. This suggests an early stage of bacterial control by the AO microenvironment followed by bacterial adaptation and proliferation. H37RvΔESX-1 presented a similar pattern of bacterial growth compared to H37Rv (Figure 1F), demonstrating that Mtb replicates in AOs irrespective of ESX-1 expression.

To determine whether AOs mounted an inflammatory/antimicrobial response to Mtb infection, we performed RT-qPCR analysis of Mtb-infected AOs 48 h post-injection focusing on cytokine, antimicrobial peptide and mucin expression (Figure 2A). Significantly induced genes included the expected IL-8 cytokine (Figure 2B)-important for immune cell chemo-attraction *in vivo*. Enhanced IL-8 secretion in the culture medium of H37Rv and H37Rv-ΔESX-1-infected organoids was confirmed by ELISA (Figure 2C). The antimicrobial peptide β-defensin-1 was also significantly enhanced upon Mtb H37Rv and H37Rv-ΔESX-1 infection (Figure 2B), which might participate in Mtb restriction during early infection. Interestingly, both Mtb H37Rv and H37Rv-ΔESX-1 significantly downregulated the expression of mucins, including MUC5B and MUC4 (Figure 2B). Mucin expression and secretion are normally enhanced during inflammation, and form part of an efficient clearance system for pathogen removal from the airway (17). Downregulation of mucin expression upon Mtb infection might facilitate bacilli transit through the airway to reach alveolar macrophages to establish its replicative niche. For all tested genes, no significant difference was observed between Mtb H37Rv and H37Rv-ΔESX-1. The H37Rv-ΔESX-1 mutant seems to induce slightly higher expression of antimicrobial peptides β-defensin-1 and −2, cathelicidin and RNAse-7, but this difference was not statistically significant (Figure 2A).

**Figure 2.**
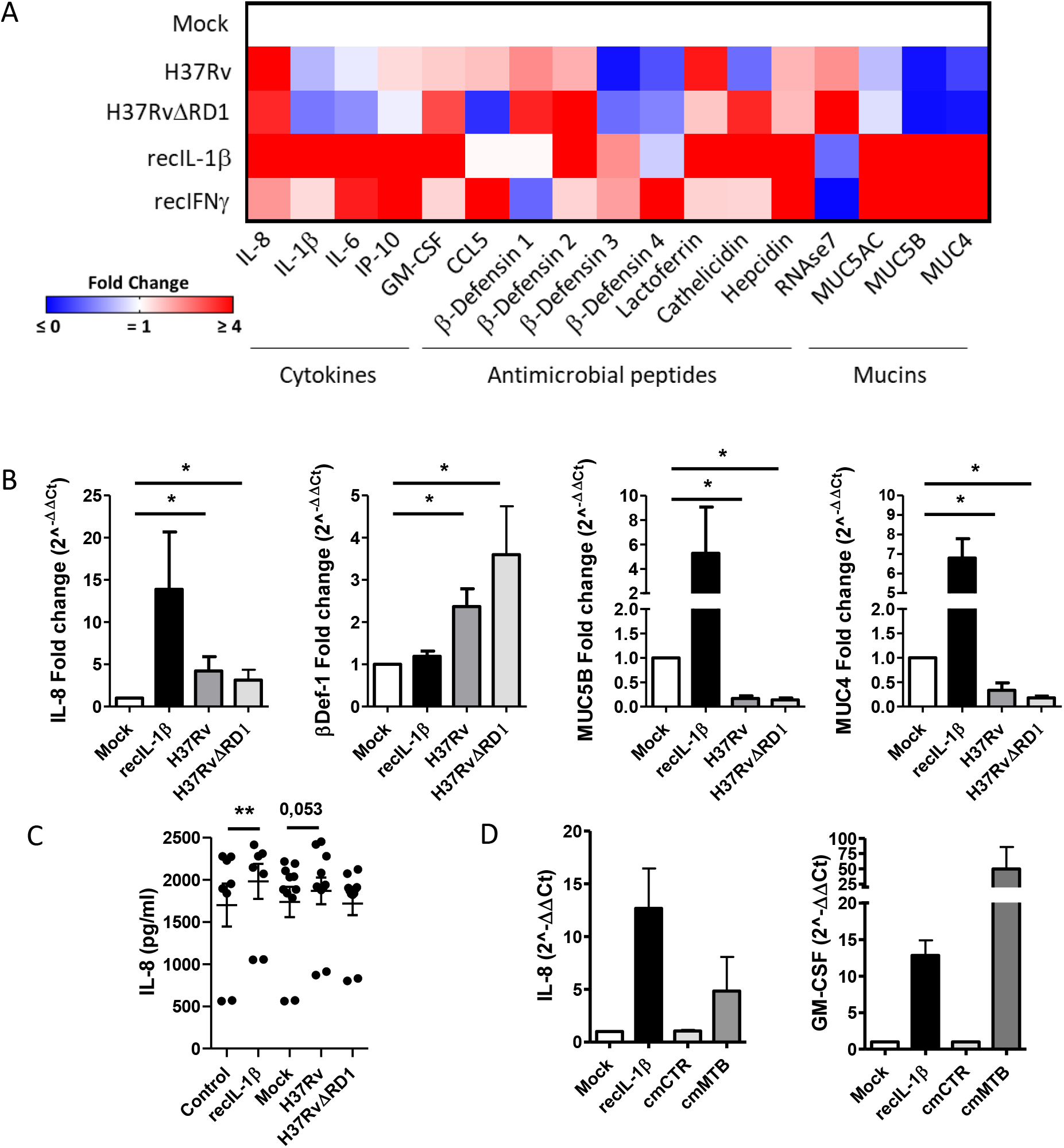
Mtb-induced host responses in AOs. (A) Heat map displaying modulation of cytokines, antimicrobial peptides and mucins in AOs in response to mycobacterial injection (H37Rv and H37RvΔRD1) compared to mock-injected organoids. As positive controls, AOs were treated with reconbimant IL-1β and IFNβ. (B) Statistically significant expression changes of IL-8, β-Defensin-1, MUC5B and MUC4 as determined by RT-qPCR at 48 h post-infection. *p < 0.05 by Wilcoxon matched-pairs signed rank test. (C) ELISA quantification of IL-8 secretion by H37Rv- or H37Rv-ΔRD1-infected AOs at 48 h post-infection. IL-8 secretion in H37Rv-infected AOs was almost significantly (*p*=0.053 by two-tailed Wilcoxon matched-pairs signed rank test), recIL-1β-treated AOs (recIL-1β) was used as positive control. (D) Statistically significant expression changes of IL-8 and GM-CSF as determined by RT-qPCR at 72 h after conditioning with cmCTR and cmMTB, defined as conditioned media from non-infected and Mtb-infected macrophages, respectively.

Upon Mtb infection, macrophages mount an inflammatory response modulating the lung microenvironment (18). AOs were stimulated with the supernatant of Mtb-infected human macrophages (cmMTB) and analyzed for gene expression compared to stimulation with the supernatant of non-infected macrophages (cmCTR). As shown in Figure 2D, among all the tested genes, the expression of IL-8 and GM-CSF, major cytokines for macrophage control of Mtb infection, were significantly enhanced in cmMTB-stimulated AOs compared to those treated with cmCTR, mimicking the paracrine macrophage-epithelial signaling occurring during lung Mtb infection. Finally, due to the essentiality of macrophages in TB disease (19), we co-cultured human monocyte-derived macrophages, alongside mycobacteria-injected organoids, and observed hourly by confocal microscopy over the course of 4 days. Due to the complex nature of this experiment, it was optimized and set up under BSL-2 conditions using *M. bovis* BCG. Human macrophages were found to migrate within the collagen matrix and in some instances, moved towards organoids containing mycobacteria (Movie S3). Whilst we found no evidence of macrophages being able to traverse the basal side and enter the organoid lumen to clear mycobacteria, we did observe some macrophages capturing and ingesting bacteria close to the basal edge of the organoid (Supp Figure 1B, Movie S3 & S4), resembling the natural process of macrophage migration to the site of infection and bacterial clearance.

We have shown that mycobacteria remain viable for up to 21 days within the lumen of AOs (Figure 1F) with approximately 2% of bacteria associating with epithelial cells after the first week of incubation (Figure 1D). During this timeframe, while AOs integrity remains uncompromised (Figure 1C, Supp Fig 1A, Movie S2), molecular interactions begin as early as 48 hours after injection with the upregulation of cytokines and antimicrobial peptides, and the inhibition of mucins (Figure 2A-C). Within 72 hours, innate immune cells can be recruited to the surface of infected AOs (Supp Figure 1B, Supp Movie S3, S4). Together, these data indicate that AOs can be used to study Mtb infection events such as primary interactions with the airway epithelium.

The ability to model these early timeframes in a responsive, multicellular and functionally similar system to the human airway, but without the complications, monetary and ethical restrictions of animal research, is revolutionary for the TB field. The ability to further introduce human macrophages allows functional modelling of a key cell type and its cellular network, overcoming a major limitation of organoid systems. We believe that this work forms the starting point for modelling a wide range of human respiratory pathogens, including SARS-CoV-2, in AOs.

## Methods

### Ethic statements

The collection of patient data and tissue for AO generation was performed according to the guidelines of the European Network of Research Ethics Committees following European and national law. In the Netherlands and France, the responsible accredited ethical committees reviewed and approved this study in accordance with the Medical Research Involving Human Subjects Act. Human lung tissue was provided by the Primary Lung Culture Facility (PLUC) at MUMC+, Maastricht, The Netherlands. Collection, storage, use of tissue and patient data was performed in agreement with the “Code for Proper Secondary Use of Human Tissue in the Netherlands” (http://www.fmwv.nl). The scientific board of the Maastricht Pathology Tissue Collection approved the use of materials for this study under MPTC2010-019 and formal permission was obtained from the local Medical Ethical Committee (code 2017-087). The CHU of Toulouse and CNRS approved protocol CHU 19 244 C and Ref CNRS 205782. All patients participating in this study consented to scientific use of their material; patients can withdraw their consent at any time, leading to the prompt disposal of their tissue and any derived material.

Human buffy coats were obtained from volunteers with informed consent via Sanquin (NVT0355.01) or établissement français du sang (Agreement 21PLER2017-0035AV02).

### Organoid culture

AOs were derived from lung biopsies as described (6).

### Bacterial culture and microinjection

DsRed-WT or −ΔESX-1 H37Rv Mtb strains were obtained by complementation with the pMRF plasmid containing a DsRed cassette, and were cultured in the continuous presence of 20 μg/ml of the selective antibiotic hygromycin and kanamycin respectively (20). Mtb strains and *M. bovis* BCG were grown and prepared for microinjection as described (18). Bacterial density was adjusted to OD_600_ = 1, and phenol red added at 0.005% to visualize successful microinjection (21). Injected organoids were allowed to recover for 2 h at 37 °C, individually collected and re-seeded into fresh matrix for subsequent analysis.

### Microscopy

For time-lapse imaging, injected organoids were seeded in IBIDI 4 well chambers (IBIDI) and stained with CellMask™ Green Plasma Membrane Stain (1/1000, Molecular Probes) for 30 min at 37°C. Organoids were washed and fresh medium containing TOPRO-3 Iodide (1/1000, Molecular Probes) was added. Organoids were imaged using a FEI CorrSight at Maastricht University or Andor/Olympus Spinning Disk Confocal CSU-X1 (10x Air 0,4 NA, 3,1 mm WD) at IPBS. Z-stacks were acquired every hour for the duration of experiments and data analyzed using Fiji/Image J and IMARIS.

For transmission electron microscopy (TEM), injected AO’s were fixed in 4% PFA for a minimum of 3 hours at RT prior to removal from the containment lab and embedding in epon blocks as described in (22). TEM data was collected autonomously as virtual nanoscopy slides on a 120kV FEI Tecnai Spirit T12 Electron Microscope equipped with an Eagle CCD camera (23).

### Colony forming unit (CFU) assay

4 to 6 Mtb-injected organoids were collected, washed in PBS, seeded into 24-well plates and cultured in complete AO medium for 7–21 days. At the relevant timepoint, organoids were lysed in 100 μl of 10% Triton X100 in water, serial dilutions were plated on 7H11 agar plates and cultured for 3 weeks at 37 °C.

### RT-qPCR

Uninfected control and Mtb-infected AO’s (15 per condition) were collected at 48 h post-infection, lysed in 1 ml of TRIzol Reagent (Invitrogen) and stored at −80 °C for 2 days. As positive controls, AO’s were stimulated with 0.02 μg/ml of human IL-1β Invivogen or 0.1 μg/ml of IFNγ (PeproTech) for 48 h. Total RNA was extracted using the RNeasy mini kit (Qiagen) and retrotranscribed (150 ng) using the Verso cDNA Synthesis Kit (Thermo Scientific). mRNA expression was assessed with an ABI 7500 real-time PCR system (Applied Biosystems) and the SYBRTM Select Master Mix (ThermoScientific). Relative quantification was determined using the 2^-ΔΔCt method and normalized to GAPDH. Primer sequences are provided in Table 1.

**Table 1.**
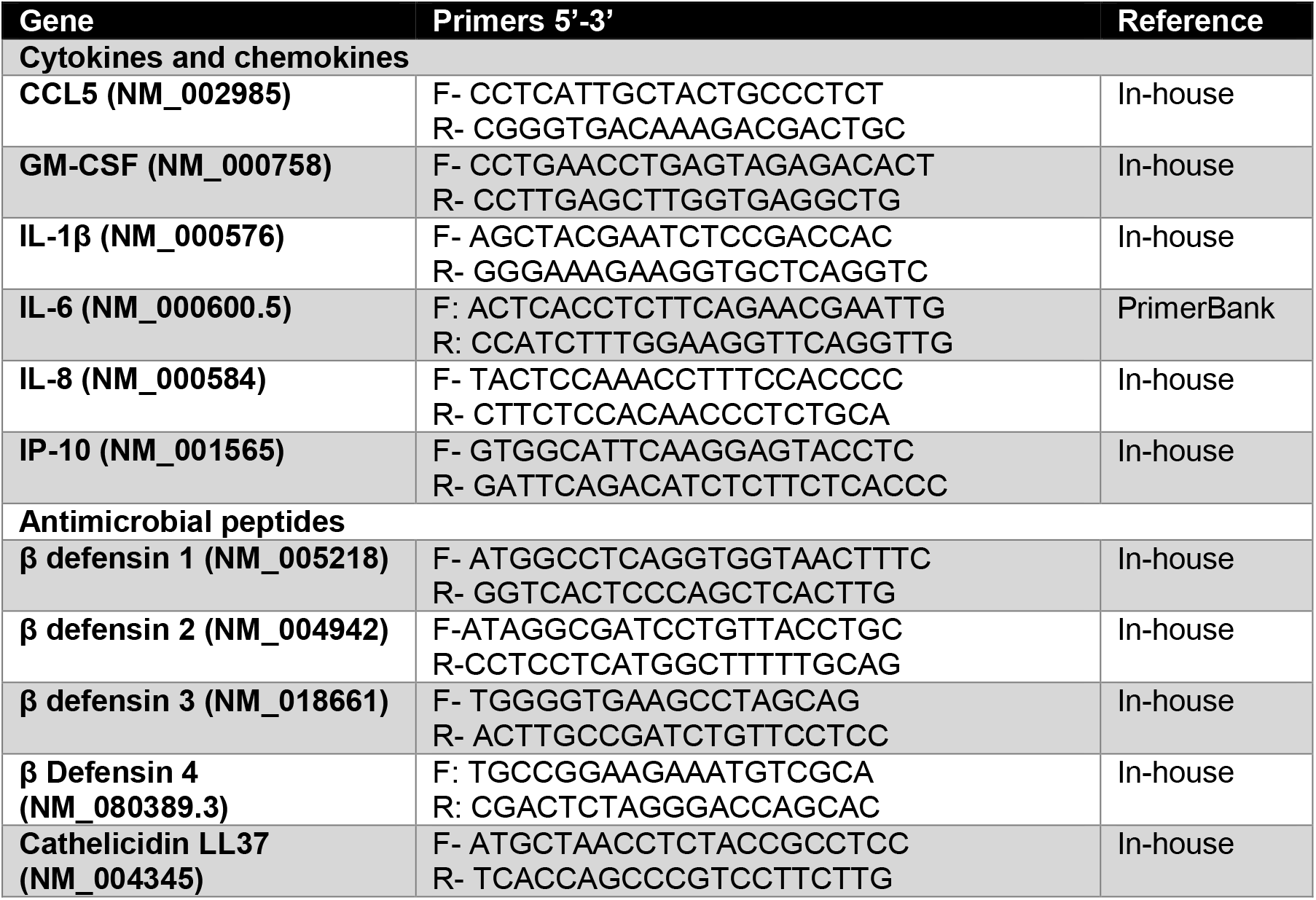
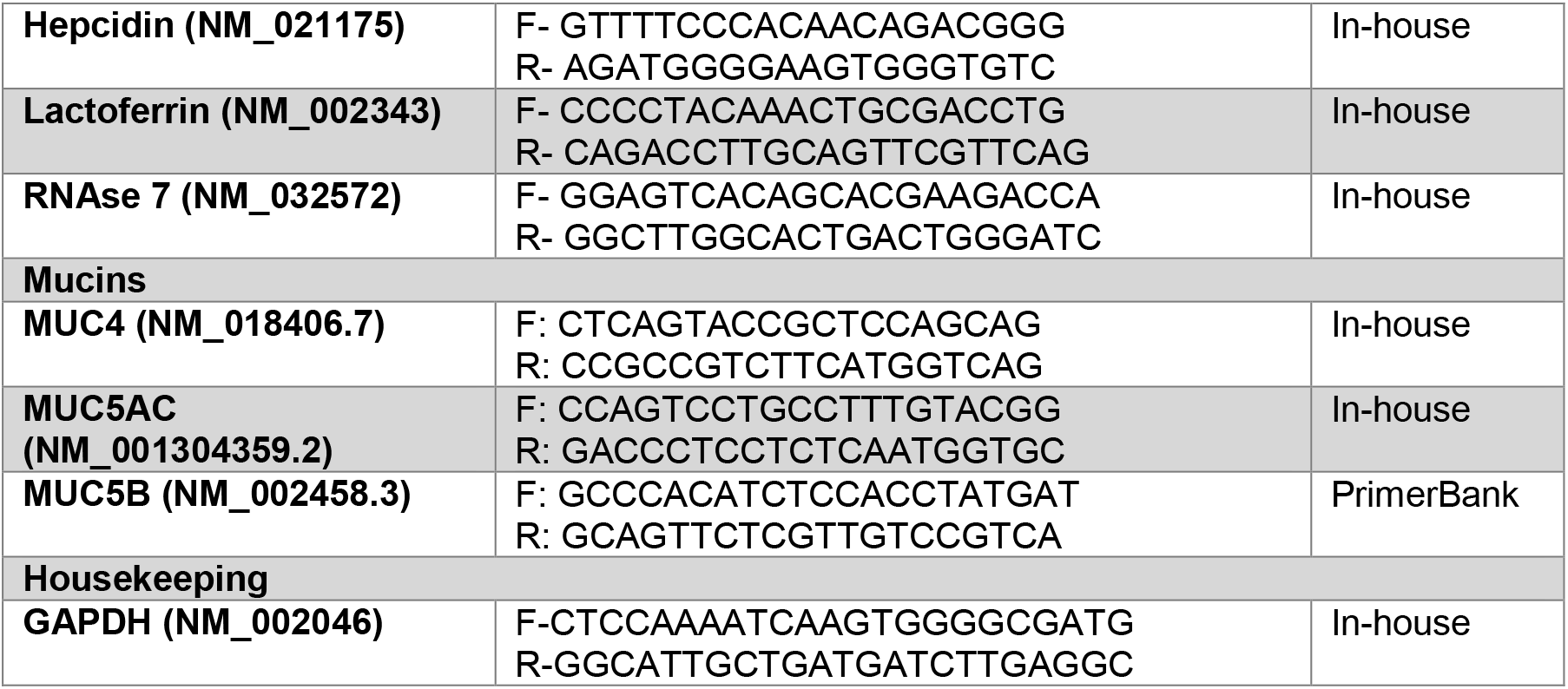
List of primers used for RT-qPCR experiments on airway organoids.

### Enzyme-linked immunoabsorbent assay

Between 20–30 organoids were embedded in fresh BME Cultrex and cultured with 800 μL complete media. After 48 h, supernatant was collected, sterilized through double 0.22 μm filters and stored at −80 °C until analysis. IL-8 ELISA was performed according to manufacturer instructions (Qiagen).

### Flow cytometry and cell sorting

Organoids were washed out of Matrigel and dissociated into single cells using TrypLE for 5 min at 37 °C. A minimum of 5 × 10^5^ cells/ml were incubated with Mtb at an MOI of 10 in complete organoid media. After 4 or 24 h for single cells, or 7 days for whole intact organoids, samples were washed with PBS, stained with CellMask Deep Red (1:30.000) and fixed in 4% paraformaldehyde overnight at 4 °C. Cells were pelleted and resuspended in PBS supplemented with 2% FCS. Samples were filtered just before analysis and sorted using a BD FACS ARIA Fusion.

### CmMTB preparation and Macrophage co-cultures

Monocytes were enriched using RosetteSep human monocyte enrichment cocktail (Stem Cell Technologies) and purified by density gradient centrifugation. Monocytes were differentiated into macrophages by addition of 5 ng/ml macrophage colony stimulating factor (Sigma Aldrich) for 6 days. cmCTR and cmMTB were prepared and used as previously described (18). Organoids were stained with CellMask Deep Red plasma membrane dye as previously described, and macrophages stained with 20 μM CellTracker Blue CMAC dye (ThermoFisher Scientific) for 1 h in serum-free media. Microinjected organoids and macrophages were resuspended in freshly prepared Rat Tail Collagen type 1 (Thermofisher, 1 mg/ml) and polymerized in a 4-well, glass-bottom μ-slide (Ibidi) at 37 °C for 30 min, and imaged for 96 h under a FEI CorrSight microscope.

## Acknowledgments

Authors acknowledge C. Kuo (Stanford University, USA) for the stable expressing Rspo1-Fc cell line and the Hubrecht Institute for the stable expressing Noggin cell line; Genotoul TRI-IPBS core facility for flow cytometry and imaging, in particular Näser and E. Vega; IPBS BSL3 facilities, in particular C. Verollet for technical support. Authors also acknowledge the Microscopy CORE lab and PLUC facility at Maastricht University. This work was supported by grants from Campus France PHC Van Gogh (40577ZE to GL-V), the Agence Nationale de la Recherche (ANR-15-CE15-0012 (MMI-TB)) to GL-V, FRM “Amorçage Jeunes Equipes” (AJE20151034460 to EM), ERC StG 693 (INFLAME 804249 to EM), ATIP to EM, ZonMW 3R’s (114021005) to PJP, the Nuffic Van Gogh Programme (VGP.17/10 to NI), and by the LINK program from the Province of Limburg, the Netherlands..

---

**Supplementary Figure 1.**
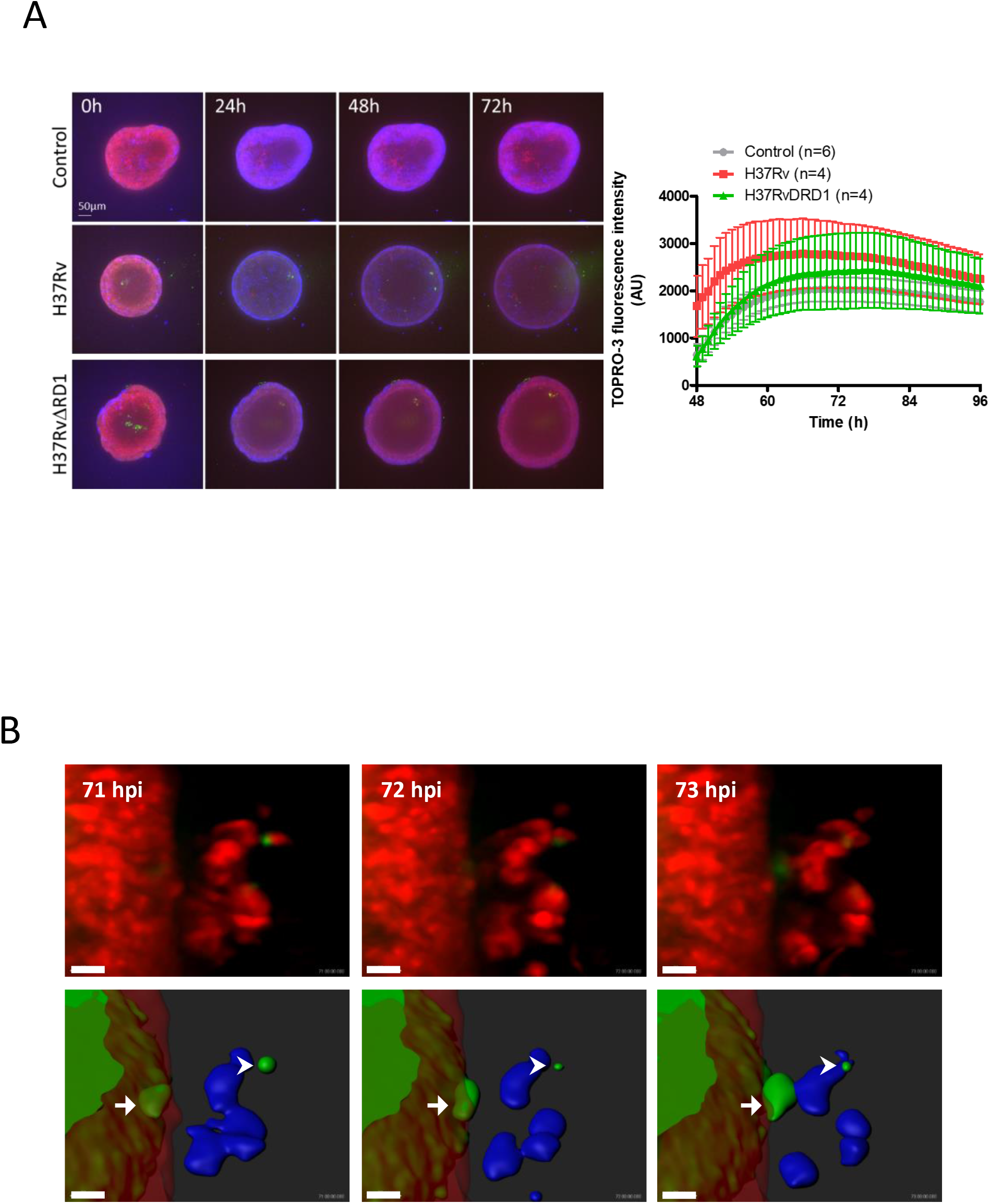
Cell death and macrophage recruitment in mycobacteria-infected AOs. Related to Figure 1. (A) AOs (red) were injected with PBS (as a control) or mycobacteria (green), stained with ToPRO3 (blue) and imaged for 4 days. ToPRO3 incorporation, and therefore epithelial cell death, was quantified using Fiji and plotted on the right panel. (B) AOs microinjected with GFP-expressing *M. bovis* BCG were embedded with human monocyte-derived macrophages in collagen and imaged hourly by confocal microscopy for 5 days. AOs and macrophages were stained with CellMask Deep Red (top row) whilst macrophages were weakly stained using Cell tracker CMAC blue allowing for segmentation in IMARIS (bottom row), arrows indicate bacteria (green) passing through the epithelial cell wall (red) and interacting with macriohages (blue). Scale bar = 20μm.

**Supplementary Movie S1. Injected mycobacteria survive in the organoid lumen. Related to Figure 1.** 3D reconstruction of an Mtb-infected AOs 4 days post-infection. DsRed-expressing bacteria are visible in red, epithelial cell membranes are stained with Cell Mask Green, and nuclei with DAPI (blue).

**Supplementary Movie S2. AO infection with WT and ΔRD1 H37Rv Mtb strains. Related to Figure 1.** Time-lapse microscopy of PBS-(left panel), H37Rv-(middle panel) and ΔRD1 H37Rv-(right panel) injected organoids (stained with Cell Mask Green) over 48 hours.

**Supplementary Movie S3. Macrophages are recruited to AOs for bacterial clearance. Related to Figure 2 and supplementary Figure 1.** Macrophages migrating to AOs in brightfield (left) and confocal microscopy (right). AOs membranes are stained with CellMask Deep Red, mycobacteria are expressing GFP and macrophages are stained with CellTracker CMAC blue.

**Supplementary Movie S4. 3D reconstruction showing frame wise interaction of macrophages with the AO surface and internal mycobacteria. Related to Figure 2 and supplementary Figure 1.** IMARIS rendering of supplementary movie S3 showing macrophages (blue) migrating to organoids (red) and cleaning up bacteria (green) from within the organoid.

## Notes

### Competing Interest Statement

H.C and N.S are inventors on patents related to organoid technology.

